# A Simple Bias Reduction Algorithm for RNA Sequencing Datasets

**DOI:** 10.1101/2023.10.31.564992

**Authors:** Christopher Thron, Hannah Bergom, Ella Boytim, Mienie Roberts, Justin Hwang, Farhad Jafari

**Affiliations:** Department of Science and Mathematics, Texas A&M, University-Central Texas; University of Minnesota

**Keywords:** RSEM (RNA Sequence by Expectation Maximization), TPM (Transcripts Per Million), FPKM (Fragments Per Kilobase of exon per Million), Local Leveling, PCA, ROC Curves, Populations

## Abstract

RNA sequencing (RNA-seq) is the conventional genome-scale approach used to capture the expression levels of all detectable genes in a biological sample. This is now regularly used in the clinical diagnostic space for cancer patients. While the information gained is intended to impact treatment decisions, numerous technical and quality issues remain. This includes inaccuracies in the dissemination of gene-gene relationships. For such reasons, clinical decisions are still mostly driven by DNA biomarkers, such as gene mutations or fusions. In this study, we aimed to correct for systemic bias based on RNA-sequencing platforms in order to improve our understanding of the gene-gene relationships. To do so, we examined standard pre-processed RNA-seq datasets obtained from three studies conducted by two consortium efforts including The Cancer Genome Atlas (TCGA) and Stand Up 2 Cancer (SU2C). We particularly examined the TCGA Bladder Cancer (n = 408) and Prostate Cancer (n = 498) studies as well as the SU2C Prostate Cancer study (n = 208). Using various statistical tests, we detected expression-level dependent, per-sample biases in all datasets. Using simulations, we show that these biases corrupt the results of *t*-tests designed to identify expression level differences between subpopulations. Importantly, these biases introduce large errors into estimates of gene-gene correlations. To mitigate these biases, we introduce *Local Leveling* as a novel mathematical approach that transforms count level data and corrects these observed biases. Local Leveling specifically corrects for the bias due to the inherent differential detection of transcripts that is driven by differential expression levels. Based on standard forms of count data (Raw counts, transcripts per million, fragments per kilobase of exon per million), we demonstrate that local leveling effectively removes the observed per-sample biases, and improves the accuracy in simulated statistical tests. Importantly, this led to systemic changes of gene-gene relationships when examining the correlation of key oncogenes, such as the Androgen Receptor, with all other detectable genes. Altogether, Local Leveling improves our capacity towards understanding gene-gene relationships, which may lead to novel ways to utilize the information derived from clinical tests.

## 1 Introduction

Gathering gene expression data from clinical samples is no longer a technical hurdle, and RNA-seq data is now frequently conducted on biological samples from the clinical and laboratory settings. When considering the holistic profile of gene expression patterns, one can inform of perturbed activity in the cell or tumor models. This information is clinically relevant since dysregulated gene activity is what altogether drives the pathogenesis of the patient’s tumors. The recognition of collective changes in tumor samples guides pre-clinical research by identifying novel signaling mechanisms and oncogenic processes. Further, this knowledge is purposed towards the development of biomarkers and precision therapies that may altogether extend the survival of cancer patients. Altogether, there is exceptional value in gathering RNA-seq data from clinical specimens. However, quality issues remain due to sequencing platforms themselves or the downstream informatics approaches and data aggregation used to interpret such data.

In current practice, gene expression levels are estimated based on RNA sequence (RNA-Seq) data obtained from populations of patients. RNA-Seq data is generated by isolating RNA from a cell or tissue, converting it to complementary DNA (cDNA), preparing a sequencing library, and using a sequencing platform to convert to transcript counts [3]. The gene transcript raw counts for each sample are typically first corrected for fragmentation effects by dividing the raw counts for each transcript by the effective transcript length [3]. This takes into account the fact that a read can occur at multiple locations along the transcript, so the number of reads per gene will be proportional to the gene transcript length. These corrected counts are then rescaled by multiplying by an overall factor. The two most widely used rescalings are transcripts per million (TPM) and fragments per kilobase of exon per million (FPKM) [1]. In particular,TPM multiplies the corrected counts so that the resulting rescaled counts sum to 1 million. It is commonly recognized that TPM is more suitable than FPKM for cross-sample comparisons [1]. Several other rescalings have also been proposed [6]. Note that the only difference between these different rescalings is an overall multiplicative factor, which varies from sample to sample.

After rescaling, additional data transforms may be applied to normalize and uniformize the data. Several types of transform are used for this purpose [4]. In particular, the log transform is widely used. The log transform data has the advantage of being easy to interpret, because differences between log-transformed values are roughly proportional to percentage differences in the original data. Altogether, each of these pre-processing steps and normalization approaches has merits and are broadly applied to datasets from consortium efforts as well as clinical reports that include RNA-seq data.

For research purposes, the pre-processed data is next used to study the tumor biology based on patterns of gene expression. The approaches include conducting differential expression analyses, gene-gene correlations, hierarchical clustering of genes based on expression patterns, enrichment of gene sets, or even machine learning approaches. Researchers adapt these approaches towards identifying the genes that altogether act as drivers or vulnerabilities of the tumors. The recurrently dysregulated genes also have potential as diagnostic tools to evaluate clinical outcomes including overall survival, the development of metastatic disease, or therapy response. Naturally, the methods in which pre-processing is conducted can act as a source of bias when analyzing patterns of gene expression, which reduced the accuracy of our understanding of gene behavior.

This study was initially motivated by preliminary investigations of differential expression levels of genes between two populations of patients within the same dataset. In multiple patient cohorts that exhibited distinct types of cancer, we identified large shifts in the overall distribution of *t* statistics when comparing random populations within the cohort. This indicated that there were per-sample biases that shifted the expression levels of all or most genes. We subsequently discovered that these shifts were systematically related to mean expression level. Altogether, if left unattended, we project these per-sample biases would corrupt any biological understanding of these samples based on conventional forms of downstream analysis.

This paper is structured as follows. In Section 2 we describe the data used, the transforms used for preprocessing, and some statistical characteristics of the preprocessed data. In Section 3 we identify the bias through block averaging of sorted data and PCA of the block averaged data. The main contribution of this paper is in Section 4, which describes a data correction filtering procedure that we term local leveling (LL). We convert the correction problem to an estimation problem, which can be used to remove the bias. In Section 5 we compare results of statistical tests performed on the original and local leveled data. In Section 6 we provide a discussion and summarize the results.

## 2 Data overview

### 2.1 Datasets used

The data used in this study comes from the three datasets consisting of samples taken from three different populations of cancer patients (one sample per patient). Two of these datasets are from the Cancer Genome Atlas Program (TCGA), while the third was funded by Stand Up To Cancer (SU2C). The datasets are summarized in Table 1. Both raw counts and TPM scaled data was used as indicated in the table.

**Table 1:** Summary of datasets used in study

| Dataset designation | Size<br>(samples × genes) | Scalings |
| --- | --- | --- |
| TCGA bladder firehose | $408 \times 20216$<br>(107 female, 301 male) | raw counts,TPM |
| TCGA prostate firehose | $498 \times 20198$<br>(all male) | raw counts,TPM |
| SU2C prostate | $208 \times 19158$<br>(all male) | FPKM, TPM |

All of the tests described in the following were conducted on all datasets. Since results were similar in all cases, the figures are shown for only one case, namely TCGA bladder firehose TPM.

### 2.2 Data preprocessing

The same data preprocessing steps were performed on all datasets. First, unexpressed genes were removed. Next, outliers (defined as expression level values more than 5 standard deviations from the mean) were clipped to five standard deviations. (Outliers are particularly common for low-expressed genes, some of which record 0 expression levels for all but a handful of samples.) The choice of 5 standard deviations was justified by the fact that for normally-distributed random variables, the probability that the random variable takes a value higher than 5 standard deviations above the mean is about 3*E* − 7, meaning that only about 3 outliers would be expected in 10 million normally distributed samples.

Next, the data was log transformed as described in Section 1, using the transformation function:

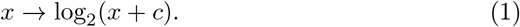

Logging the data compresses the range (expression levels typically range from close to 0 up to close to 100,000, with most between 1 and 100) and facilitates comparison between genes that express at very different levels. Furthermore, the distributions of log transformed expression level data for individual genes are more nearly normal than the original untransformed data.

The constant *c* is included in (1) so that unexpressed genes will not transform to negative infinity. Some investigators take *c* = 1, but this distorts the scale for low-expressed genes. Taking *c <* 1 may reduce this distortion, but if *c* is taken too small then variance for the log transform of low expressed genes becomes inflated. For most datasets we chose *c* = 0.25, but for FPKM data we chose *c* = 0.125 because expression levels tend to be lower, due to the different normalization.

In many investigations, the per-gene expression levels is converted to standard scores (*Z* scores), so that the per-gene expression levels all have mean 0 and standard deviation 1. In our case we did not standardize because the bias’ effect on a given gene depends on its mean expression level, but not on its standard deviation. Dividing each gene’s expression level by its own standard deviation will scramble the bias so that it can no longer be estimated or corrected. (Note however that the statistical tests in Section 5 do apply standardization to compute *t* statistics, but only *after* bias correction.)

### 2.3 Statistical characteristics of preprocessed data

In this section we establish some normality and uniformity properties of log transformed RNA-seq data. We find that gene averages of small populations are nearly normal, and that the log transform tends to uniformize the variance across the scale of expression levels.

#### 2.3.1 Normality of per-gene expression data

Many statistical tests depend on the assumption of normality. In order to characterize the normality of log transformed expression level data, we applied the d’Agostino test for normality to the transformed data for each gene across samples. Figure 1 shows the sorted d’Agostino *p*-values for all genes for the original data. A straight line from (0,0) to (1,1) is consistent with perfect normality–however, we find that for the original data, most *p*-values are 0, indicating poor agreement with normality.

**Figure 1.**
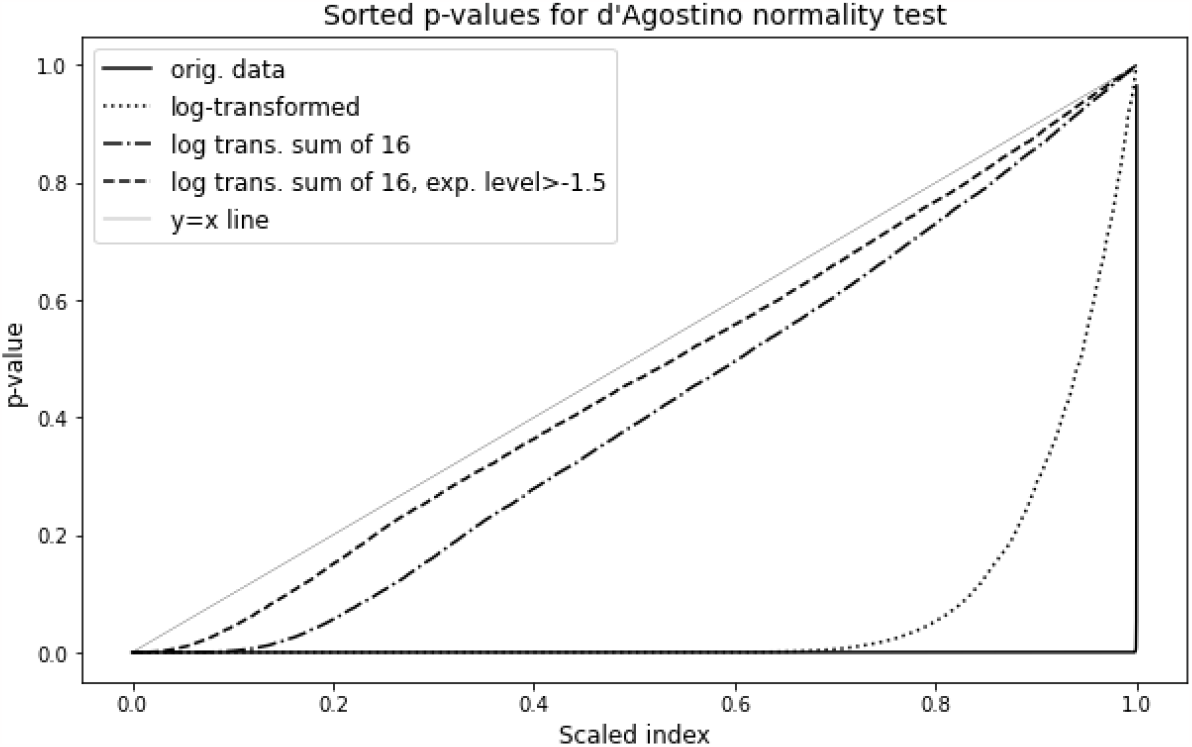
Sorted *p*-values for d’Agostino test applied to per-gene data across samples for untransformed data, log transformed data, log transformed sum of blocks of 16 patients, and log transformed sum of blocks of 16 patients for genes with log transformed expression levels above 1.

The figure also shows sorted d’Agostino *p*-values for log transformed gene data. The normality is somewhat improved, so that about 20 percent of the genes have *p* values greater than 0.05. The normality is further improved if we look instead at the distribution of per-gene averages across patients. Figure 1 also shows *p*-values obtained from averages of 16 samples from the preprocessed TGCA bladder cancer dataset. Since this dataset has about 400 samples, this means that each data point is calculated based on the distribution of 25 values. Most of the curve is linear, indicating greater normality. However, there are still about 10% of the genes that have *p*-values indistinguishable from 0 on the scale of the figure. Examination of the data reveals that very low expressed genes have highly irregular distributions. It follows that low-expressed genes are likely to exhibit non-normality. Accordingly, we selected only those genes with mean transformed expression level greater than -1.5 (about 83% of all genes), and plotted the sorted *p*-values for averages of 16 for this restricted set of genes. This time, the *p*-value distribution is very close to what would be expected from normally-distributed data. This result justifies the assumption of normality in tests that compare mean expression levels in subpopulations, as long as the subpopulation size is greater than 16.

#### 2.3.2 Uniformity of per-gene expression data

We also examine the expression level dependence of per-gene standard deviations for log transformed data. Figure 2 shows the standard deviations of log transformed gene expression levels for the bladder cancer TPM dataset. In the plot, genes are ordered in increasing order of mean expression level. The plot shows that the standard deviations of log transformed data are indeed nearly independent of expression level. The color bands in the plot indicate the ranges for the five data quintiles: for example, about 60% of transformed expression levels are between -1 and 5. The near-uniformity of per-gene standard deviations across all expression levels shows that per-sample expression level deviations from the mean are the same order of magnitude across all genes, regardless of expression level. We will see that the effect of expression-level dependent bias is also of the same order of magnitude across all expression levels.

**Figure 2.**
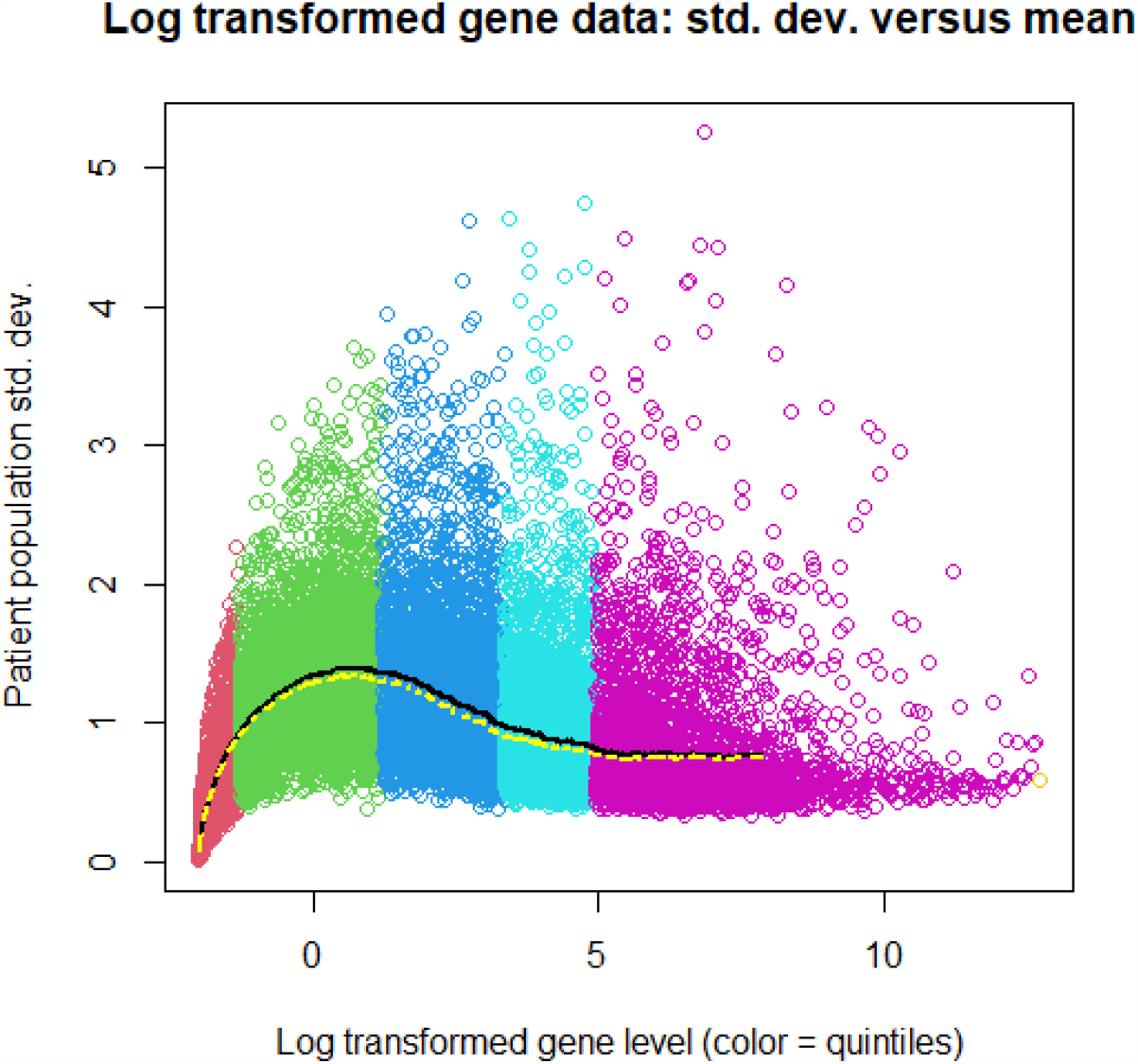
Standard deviations of log transformed genes sorted by mean of transformed expression level. The color bands indicate quintiles of log expression levels from 0-20%, 20-40%, 40-60%, 60-80%, and 80-100%. The black line is the moving average of standard deviations, where each point on the line is the average of 1000 consecutive genes. The yellow dashed line is the moving average of standard deviations after local leveling.

## 3 Bias detection and characterization

In this section we describe various statistical tests that detect per-sample expression level biases in the studied datasets, and show that they are related to gene expression level.

### 3.1 Randomized 2-population *t*-tests

The first test is designed to show the sensitivity of two-population *t* tests to the presence of actual genetic differences between the two populations. For this purpose, we conducted 5000 trial simulations. In each trial we randomly chose a subpopulation of 107 samples, computed 2-population *t* statistics for each gene, and recorded the top 2000 *t* values (out of 20216 total genes). For each of these 5000 cases, for a range of *t* levels we recorded the number of genes exceeding each *t*-level. The resulting 5% and 95% confidence limits for each *t* level are plotted as solid red curves in Figure 3. These results show an enormous variation in *t*-statistic distributions among random population divisions. For example, for *t* = 3 the [5%,95%] confidence interval is roughly [0, 1000], meaning that different choices of random subpopulations may produce as few as 0 and as many as 1000 gene outliers with *t* statistic greater than 3.

**Figure 3.**
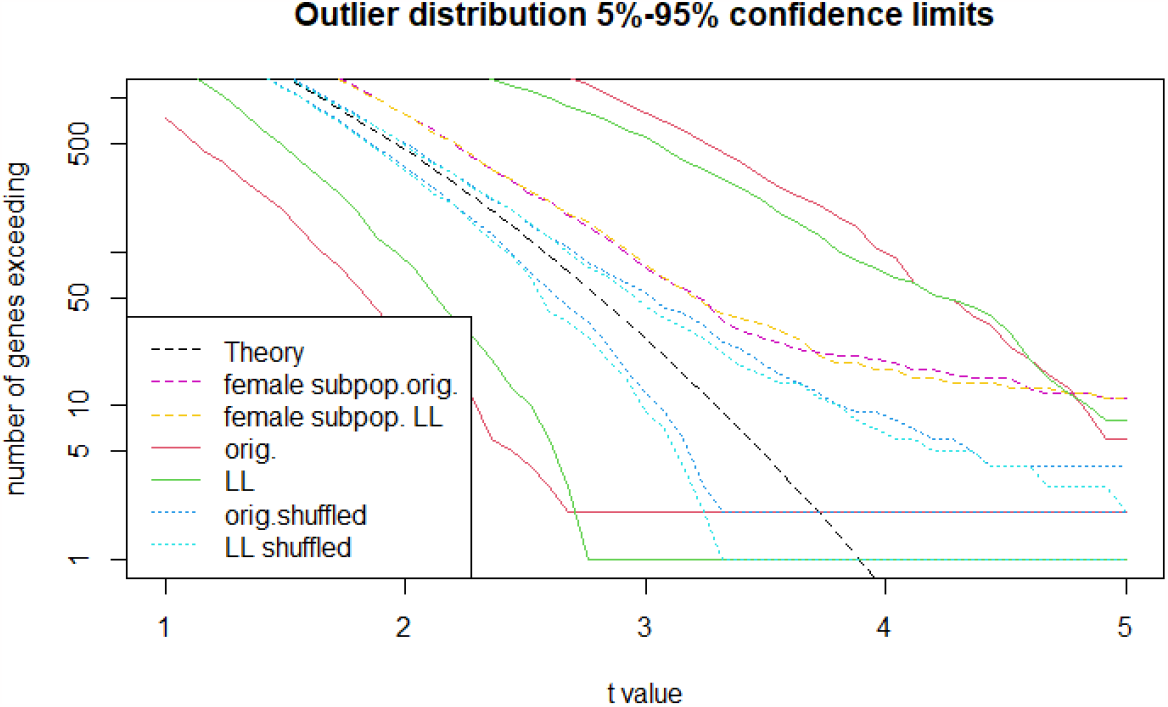
Confidence limits on complementary cumulative distribution functions for *t*-tests between randomized subpopulations. Dashed black line indicates theoretical *Z* distribution assuming no statistically significant differences between populations. the two red solid lines show the 5% and 95% confidence limits for original log-transformed data. The red dashed line shows the case when the subpopulation consists of all female samples. Dashed blue lines show 5% and 95% confidence limits when the original log transformed data is shuffled gene by gene. The solid green curves display the 5% and 95% confidence limits after local leveling, and the dashed cyan lines show the shuffled local leveled case.

On the other hand, if the gene expression levels are first shuffled gene-by-gene among patients and then the same procedure is followed, the confidence limits on *t*-statistics are much tighter (blue dotted lines in Figure 3), and approach the theoretical *t* distribution (black dashed line in Figure 3). The comparison between shuffled and unshuffled samples show that there are shifts in expression level from sample to sample, which affect multiple genes in each sample. When the expression levels are first shuffled among samples, then genes from the same sample are no longer together, so the overall shift is no longer observed.

We may compare these results with a case where subpopulations have known genetic differences–in this case male versus female. The dashed red curve in Figure 3 shows the gene outlier curves for the subpopulation consisting of all 107 female samples in the dataset. In this case, when *t <* 4, the number of genes greater than *t* is well within the confidence limits. It follows that the large variation in outlier distributions are not due to sex differences. Only *t* values larger than 5 can definitively identify genes as sex-linked. If there are other genes that tend to be elevated in females, then they cannot be clearly identifiable based on *t* values alone.

The observed variation in outliers could possibly be due to a few samples that are skewing the results. We may test this hypothesis by correlating patients’ inclusion within the subpopulation with number of outliers. The red curve in Figure 4 shows the sorted correlations of patient inclusion in the subpopulation (a 0-1 random variable) with the number of genes having *t* value greater than 2.5. This may be compared with the blue dashed curve in the same figure, which shows sorted correlations generated similarly except with per-gene expression levels shuffled among patients. We see that a large number of samples have raised (or lowered) correlations, indicating that the variation in outliers cannot be attributed to just a few patients, but rather most samples will have either a net positive or net negative effect on the number of outliers if included in the subpopulation.

**Figure 4.**
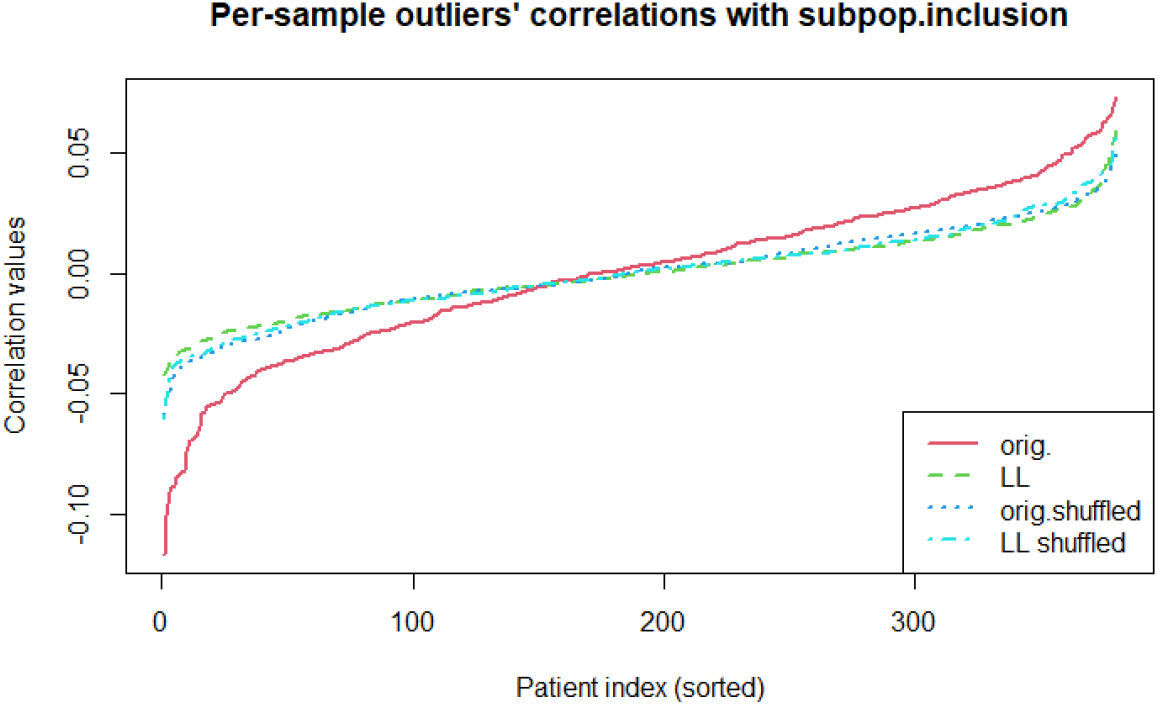
Correlation of sample inclusion within subpopulation and number of gene outliers with *t*>2.5, for the 2-population tests described in Section 3.1. Curves for tests with original log transformed data (red), Local leveled (green dashed), original log transformed data shuffled by gene (blue dashed), and local leveled shuffled (cyan dashed) are shown.

We may similarly test the hypothesis that there is a limited subset of genes that are involved in producing the large variation in the number of outliers observed in Figures 3. The red curve in Figure 5 shows sorted correlations of the mean expression levels of genes included in the subpopulation with the number of genes having *t* value greater than 2.5. The blue dashed curve in the same figure shows sorted correlations generated similarly except with per-gene expression levels shuffled among patients. Evidently the majority of genes have raised correlations, indicating that the effect cannot be attributed to just a few genes.

**Figure 5.**
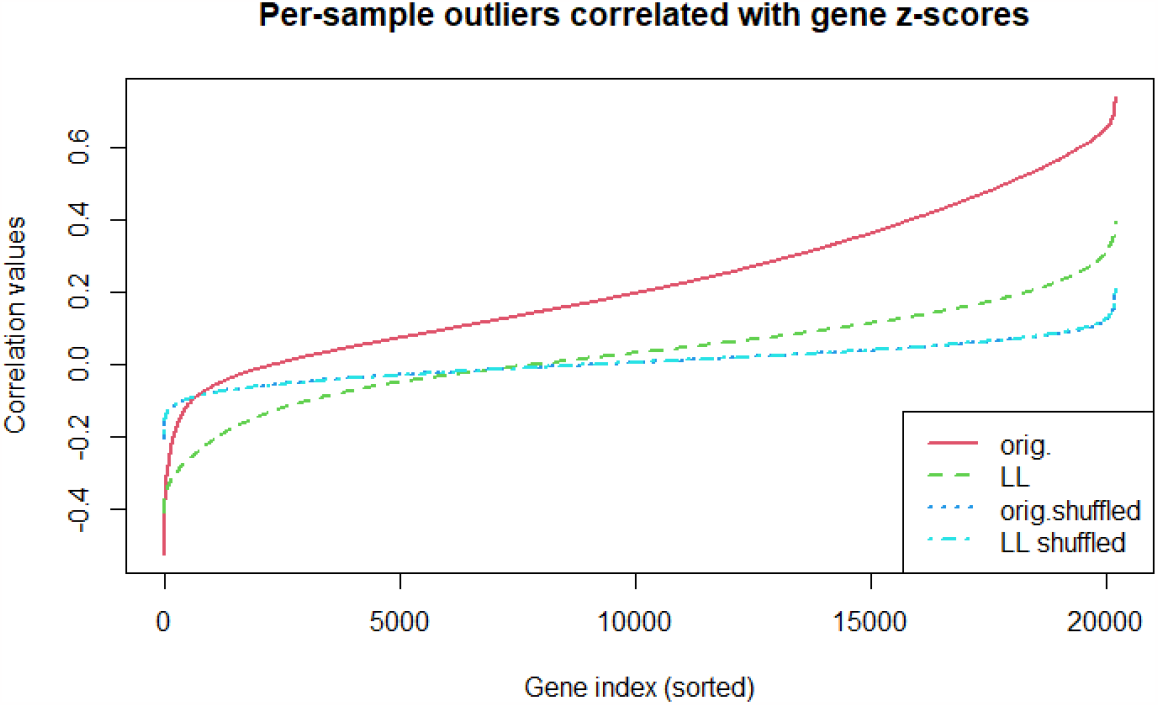
Correlation of mean expression level of genes of included samples with number of gene outliers with *t >* 2.5, for *t* tests described in Section 3.1. Curves for tests with original log transformed data (red), Local leveled (green dashed), original log transformed data shuffled by gene (blue dashed), and local leveled shuffled (cyan dashed) are shown.

### 3.2 Block averaging of sorted expression levels

In Section 3.1 we saw that there appears to be sample by sample shifts in expression level. In this section, we investigate whether the size of these shifts has any relation to the gene’s expression levels. This task is complicated by the fact that shifts produce a relatively small effect compared to random expression level variation on a per-gene basis. However, the relative effect of the shifts is amplified by taking per-sample block averages of sorted expression level data.

The procedure for detecting expression-level shifts is described in detail as follows. Note that “expression level” refers to the expression level of data that has been cleaned and log transformed as described in Section 2.2.

1. Compute the per-gene mean expression levels across all samples, and then sort the genes in order of increasing mean value.
2. Divide the sorted expression levels into consecutive blocks of equal size, and sum the transformed expression levels in each block separately for each sample. This step is made mathematically precise as follows. Let *J, K* denote the number of genes and samples respectively, and *g*_*jk*_ denotes the transformed expression level for gene *j* (*j* = 1 … *J*) and sample *k* (*k* = 1 … *K*). Let *B* denotes the block size, and let *b*_*nk*_, (*n* = 1 … *N, k* = 1 … *K*) denote the average expression level for all genes in block *n* and sample *k*. Then we have:

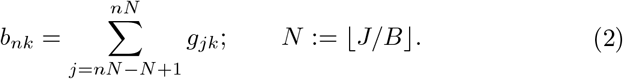
3. In order to compare across samples, we then center the values {*b*_*nk*_} by subtracting out block means across samples to obtain centered block averages 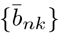 as follows:

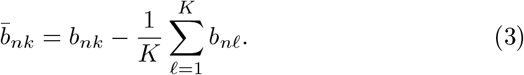

Figure 6(a) shows centered block averages as defined by (3) for selected samples from the TPM bladder cancer data, with block sizes of *B* = 1000. The trends show that individual samples have smoothly varying deviations from mean gene expression level depending on expression level.

**Figure 6.**
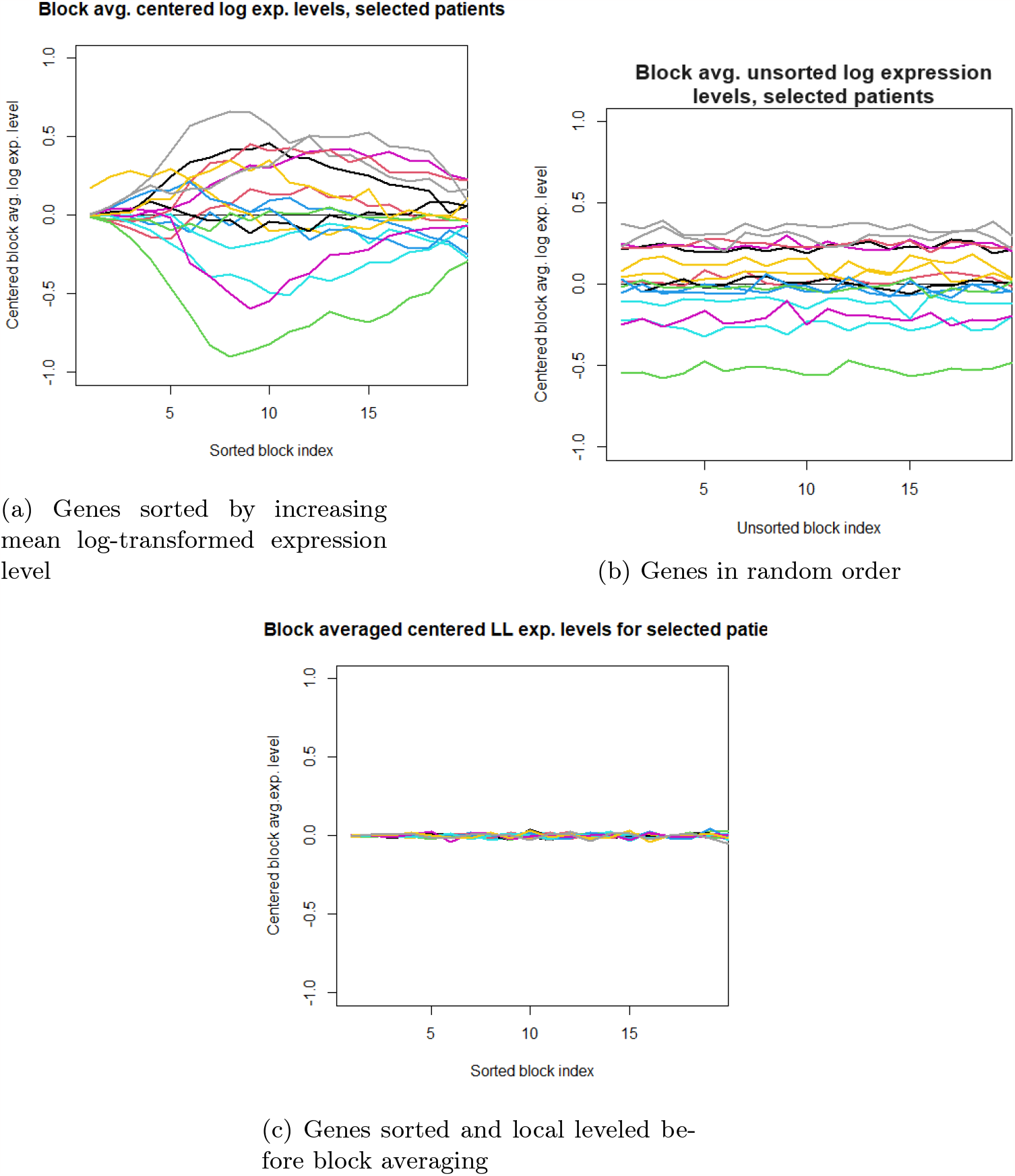
Centered block averages (B=1000) for selected samples from the TPM bladder cancer data.

In order to confirm that the deviations shown in Figure 6(a) are dependent on expression level, we repeat the process (1-3) described above, but without sorting the genes by increasing expression level. Figure 6(b) shows that block averages of unsorted genes do not show varying deviations: instead, there are constant overall shifts from sample to sample. This confirms the systematic effect of gene expression level on per-sample expression level deviations from the mean.

### 3.3 Principal component analysis of block averaged data

Principal component analysis (PCA) is a well known tool to reduce the dimensionality of high-dimensional data. We can use PCA to analyze the deviations shown in Figure 6. Figure 7(a) shows the first three principal components for the block averaged sorted data described in Section 3.2. These results may be compared to Figure 7(b), which shows the first three principal components for block averages of the same data, but unsorted. The sorted block averages show clear trends, indicating systematic expression-level dependent deviations from the mean.

**Figure 7.**
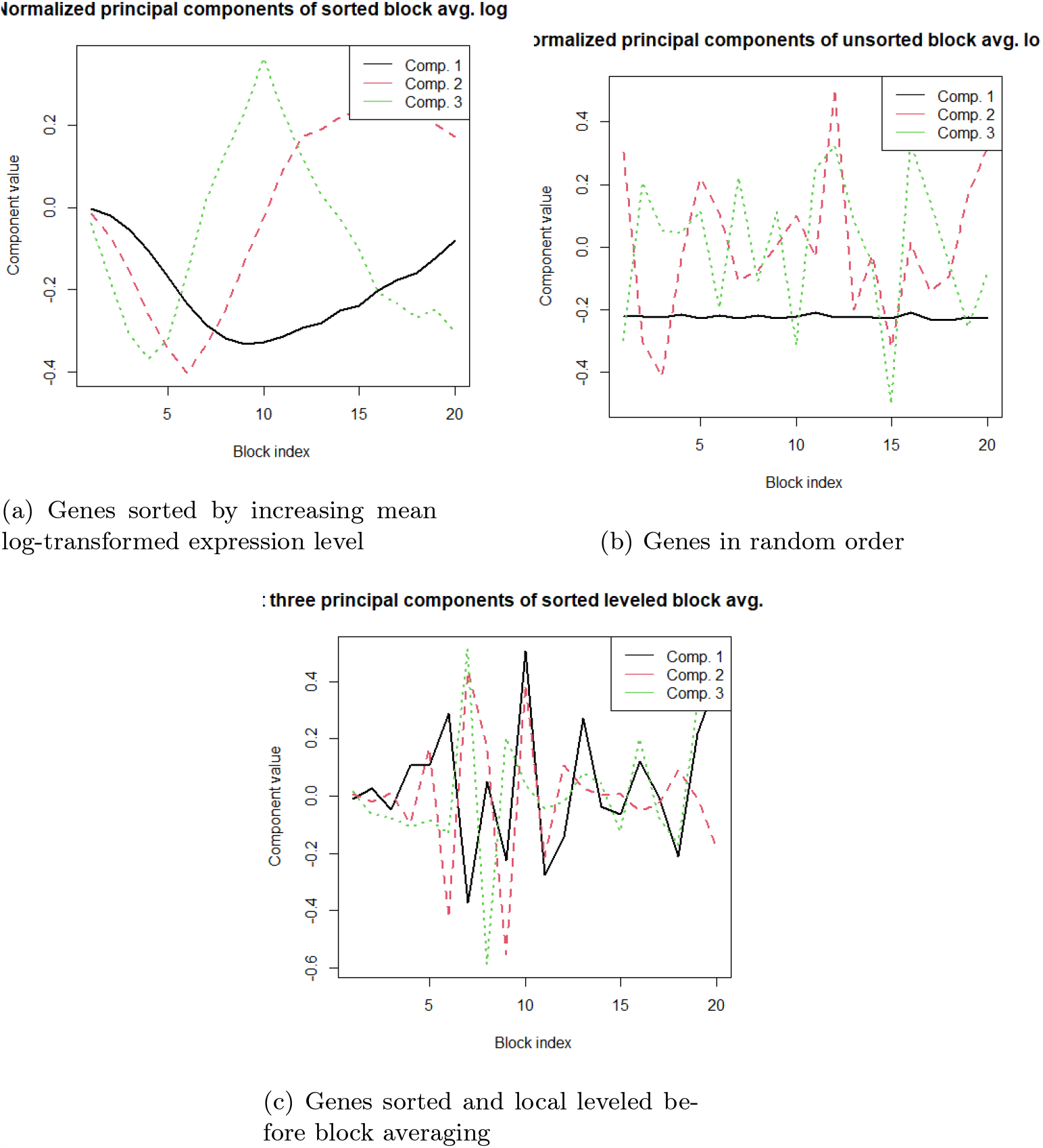
Leading PCA components for block averaged data with different orderings and with local leveling.

Figures 8(a) and (b) shows the explained variance associated with the principal components shown in Figures 7. For the block averages of sorted data, the first two components account for 95% of the total variance, implying that the per sample deviations can be almost completely characterized as linear combinations of the first principal components. On the other hand, Figure 8(b) tells a very different story for block averaged unsorted data. The block averaged per-sample deviations are uniform, signifying that the varying trends seen in sorted data have been smeared out.

**Figure 8.**
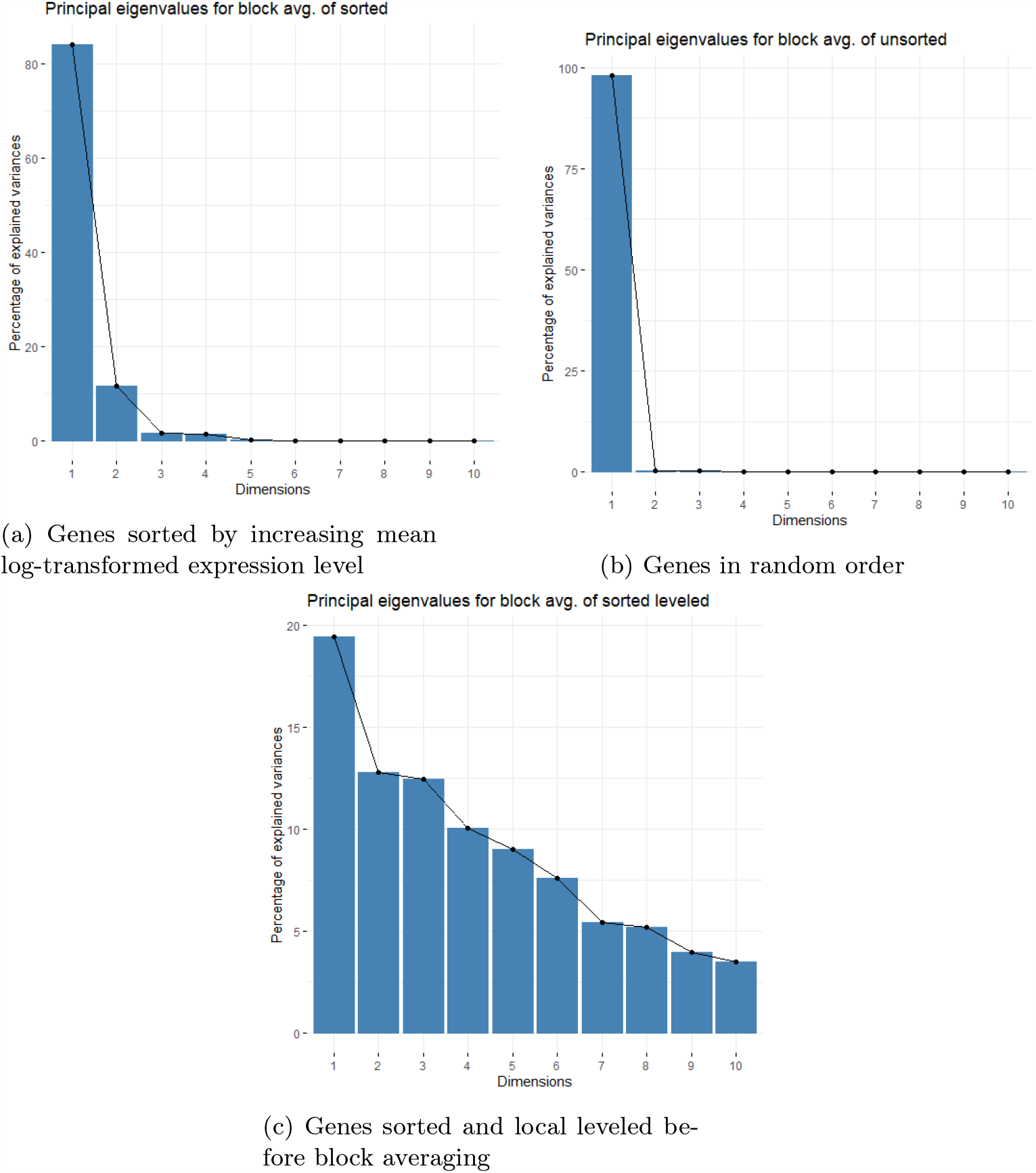
Eigenvalues corresponding to the PCA for block averaged deviations for: (a) Sorted transformed data; (b) Unsorted transformed data; (c) Local leveled sorted transformed data.

### 3.4 Implications of statistical test results

Sections 3.1-3.2 have established the existence of systematic shifts in gene expression levels that vary from sample to sample and depend on the expression levels. These shifts apparently affect very large numbers of genes. We can ask if the shifts are of biological origin, or if they are due to measurement bias. The fact that the shifts depend on expression level independent of gene function strongly indicates that these results do not have biological significance, but rather reflect measurement bias. Accordingly, in the subsequent discussion we will refer to the effect as a bias rather than a shift.

## 4 Bias correction through local leveling

In Section 3 we have shown the existence of sample-by-sample bias, and have shown that it has a large effect on t-tests such as could possibly be used in clinical studies. Undoubtedly this bias will introduce errors into the results of such tests, and may obscure effects due to varying levels of real activity among patients or groups of patients. It behooves us therefore to find a way to reduce these biases. The most straightforward way to do this is to estimate and then subtract the bias. The correction problem is then reduced to an estimation problem.

Let *X*_*p,g*_ be a random variable expressing the expression level of gene *g* in sample *p*. Then we may characterize *X*_*p,g*_ using the following probability model:

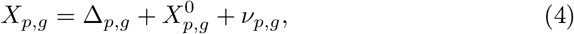

where Δ_*p,g*_ is the bias for gene *g* for sample *p*; 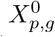 is the true (unbiased) expression level of gene *g* in sample *p*; and *ν*_*p,g*_ is mean-0 random statistical variation.

Our goal is to estimate Δ_*p,g*_. For this purpose, we will make use of the properties of statistical averaging. First we define notation: for any random variable *Y*_*p,g*_ depending on *p* and *g* and for any set of genes *𝒢*, let 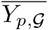 denote the average of *Y*_*p,g*_ over all genes *g* ∈ *𝒢*:

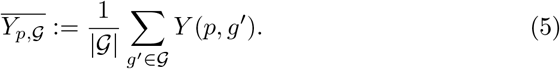

Then we have:

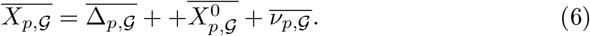

We may similarly denote the average of *Y*_*p,g*_ over genes *g* ∈ G and samples *p* ∈ P by 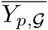 Then we have also:

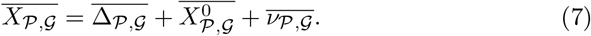

By rearranging and combining (6) and (7), we obtain:

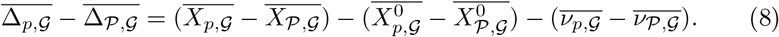

But since Δ_*p,g*_*′* is the mean bias for sample *p* and gene *g*^′^, it follows that the average over all samples of Δ_*p,g*_*′* is 0 for all genes *g*^′^. It follows that 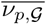, so (8) simplifies to:

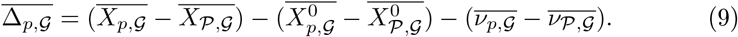

Note the first term on the right-hand side of (9) is calculated from data, while the second and third terms cannot be measured directly. To estimate the second term, we assume that most of the genes in each sample are “typical” in the sense that they do not have systematically elevated or depressed levels relative to the same genes in other samples. Under this assumption, the true average of expression levels for a large number of genes should not vary from sample to sample. In other words, we have the condition that

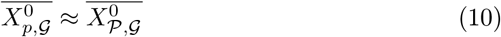

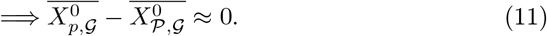

As to the third term on the right-hand side of (9), since *ν*_*p,g*_*′* has zero mean and finite variance, by the Law of Large Numbers we have that the averages 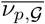 and 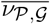 both approach 0 as increases. This fact together with (9) and (10) gives

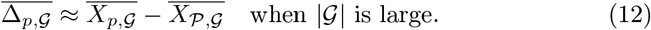

Now suppose that given a gene *g*, we can choose a set of genes *𝒢* such that 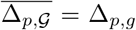. Then we can use (12) to estimate the bias Δ_*p,g*_ for person *p* for gene *g*. The method therefore comes down to choosing a relatively large set of genes *𝒢* whose average bias is equal to Δ_*p,g*_. We can see from Figure 6 that per-sample biases have approximately linear dependence on expression level for blocks of consecutive genes of size less than a few thousand. Therefore for a given *g*, if we choose *𝒢* to be a consecutive set of *B* genes where *B* ≈ 1000, such that the 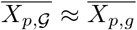 for all samples *p* ∈ *𝒫*, it follows from linearity that 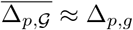 for all samples *p* ∈ *𝒫*.

In our experiments, we chose the sets {*𝒢*} as follows. First, let *g*_1_, … *gN*_*g*_ denote all genes sorted by expression level. We defined gene sets *𝒢*_*n*_, *n* = 1, … *N*_*g*_ − 1000 as follows:

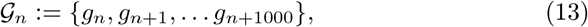

i.e. *𝒢*_*n*_ contains 1001 consecutive genes beginning at *g*_*n*_. Let *m*_*n*_ be the midpoint expression level for *𝒢*_*n*_,

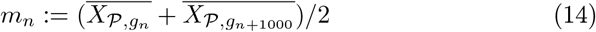

Then for each gene *g*, we assign the set *𝒢*_*n*_ \ {*g*}, where *n* is chosen such that

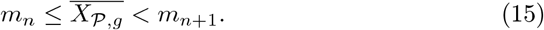

In other words, the midpoint of *𝒢*_*n*_ is the largest midpoint that is smaller than mean expression level of *g*. According to this specification, *g* is nearly equal to the midpoint of the expression level interval spanned by the genes in *𝒢*_*n*_.

We may graphically assess the effects of local leveling. In Figure 2, the yellow dashed line gives moving averages of per-gene standard deviations of local leveled log transformed data, plotted against mean expression level. The standard deviations are slightly reduced compared to the black line which represents the nonleveled data. This shows that the standard deviations of the log transformed gene expression levels are not changed significantly by local leveling.

A much larger effect of local leveling is seen in block averages of per-sample deviations. Figure 6(c) shows that block-average deviations virtually disappear after local leveling. Figure 8(c) shows that different PCA components make comparable contributions to the per-sample deviations. It appears that local leveling has effectively removed the per-sample deviations, since both the expression-level dependence seen in sorted data and the overall shifts seen in unsorted data have been removed.

## 5 Comparison of statistical tests with original and local leveled data

We have shown that the local leveling transform eliminates biases in block-averaged, sorted data. In this section we investigate the effect of local leveling on some statistical tests that may be used in clinical studies.

### 5.1 Summary of test types

Three types of tests are performed, in order to determine the effect of local leveling on detectability of genetic differences. The three types are described in the following three subsections.

#### Detection of differences in gene expression level between subpopulations

As in Section 3.1, the population is divided into two subpopulations. In one of the subpopulations, for a randomly chosen set of genes the expression levels are altered in a pre-determined way by enhancing them by a multiplicative factor. *t*-statistics are computed for all genes, which are then ranked in decreasing *t* order. Different ranks are chosen as detection thresholds, and the number of enhanced genes that are detected are recorded for both original and local leveled data. Detection rates of the enhanced genes are plotted for both original and local leveled data, for different thresholds and different enhancement factors.

#### Detection of individual samples with elevated expression levels in a known set of genes

A sample is chosen randomly, and a randomly-chosen subset of genes are enhanced by multiplying them by a pre-determined factor greater than 1. Then *t*-statistics for the genes in the subset are used to rank the samples. The rank of the sample for original and local leveled data is compared, for different subset sizes and enhancement factors.

#### Effect of leveling on correlations

The distribution of gene-gene correlations are compared for genes with similar expression levels versus randomly-selected genes. The block-averaged results shown in Figure 6 imply that genes with similar expression levels will tend to be correlated more than other genes. We compute the same correlations with original and local leveled data, to see whether local leveling removes biases in estimates of correlations between genes.

### 5.2 Results of statistical tests

#### Subpopulation comparisons

Figure 9 shows the results of the two population test performed for 5000 trials, where 100 randomly-chosen genes were enhanced in a randomly-chosen subpopulation of 100 samples. The three series of curves correspond to three different enhancement factors (1.1, 1.3, 1.5). The curves show local leveling consistently produces a slight improvement in detectability. For the larger enhancement factors the standard deviation (indicated by error bars) is also decreased when local leveled data is used, indicating that more consistent results are obtained.

**Figure 9.**
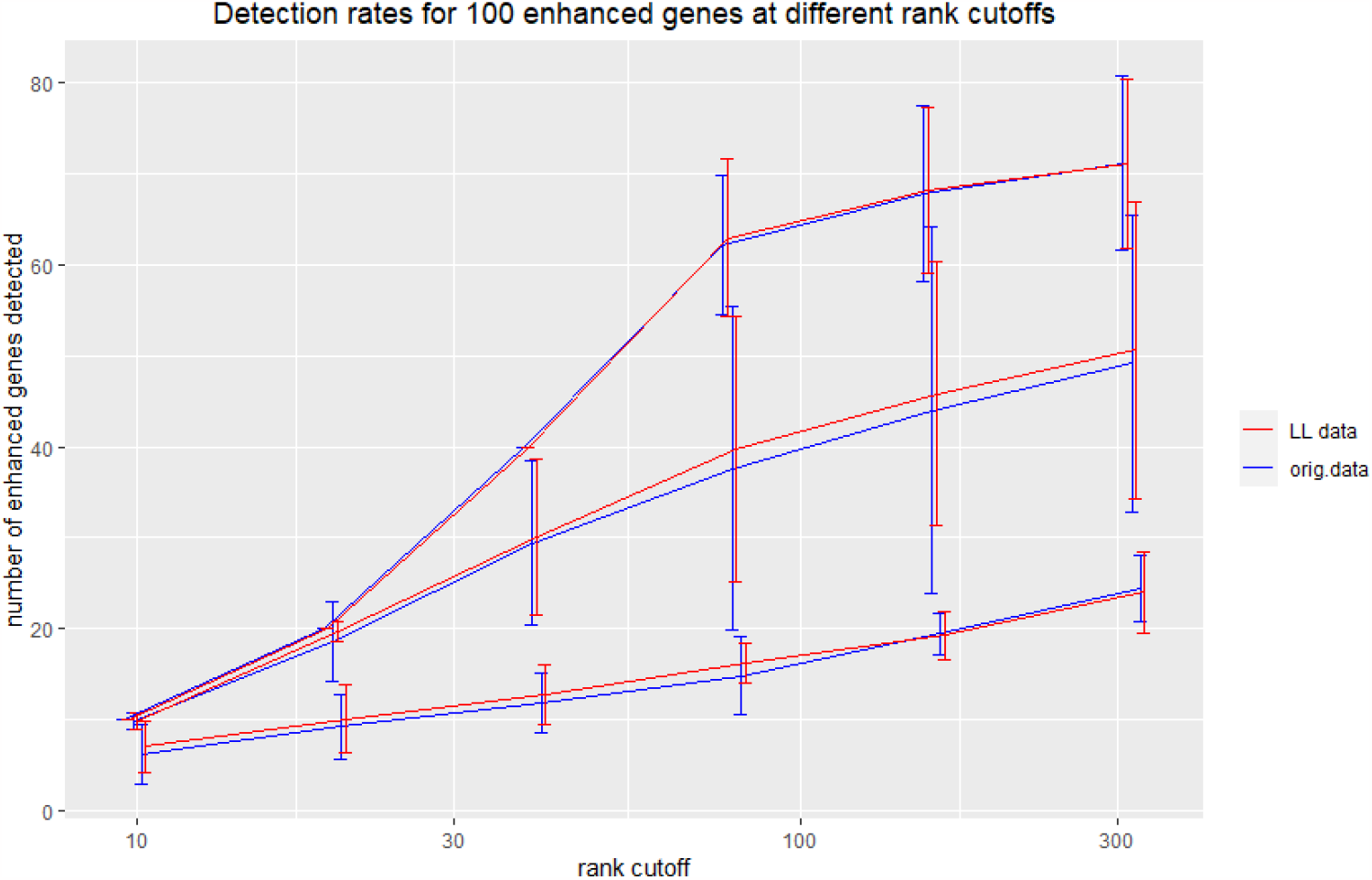
Detection rates for 100 enhanced genes in a subpopulation of 100 at different rank cutoffs, with and without local leveling. The three series of curves correspond to three different enhancement factors (1.1, 1.3, 1.5). Means and standard deviations are shown for 5000 trials with 100 randomly-chosen genes from 100 randomly-chosen samples.

#### Elevated single-sample detection

Figure 10 shows the results when the single patient test was performed for 1000 random sample/gene combinations for each enhancement factor and each enhanced genes subset size. The means and standard deviations of *t* ranks for two different enhancement levels and for original and local leveled data for a range of gene set sizes correspond to the four curves with error bars shown in the figure. Compared to the original data, the local leveled data improves the rank of the gene-enhanced sample considerably, as well as reducing the error bars. For example, with 40 enhanced genes and enhancement factor 1.4, the mean *t* rank for local leveled is 1 (with small error bars) compared to about 9 for original data (with much larger error bars). The improvement increases dramatically with an increase in the number of enhanced genes.

**Figure 10.**
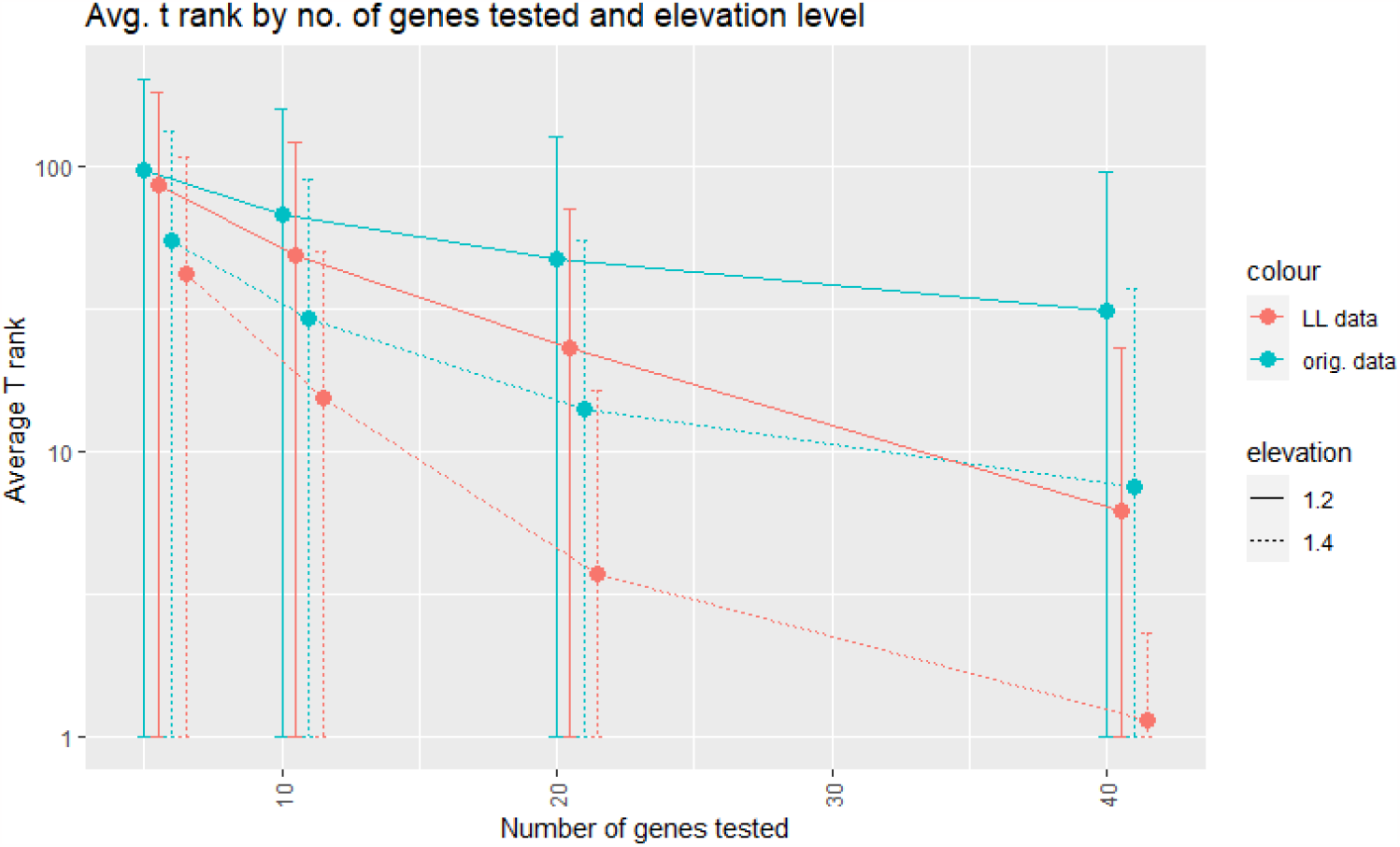
Average *t* rank for a single sample with variable number of enhanced genes. The number of enhanced genes is indicated on the *x*-axis. The solid and dotted curves correspond to two different multiplicative enhancement factors for original (blue) and local leveled (red) data. Means and standard deviations are shown for 5000 trials, where the enhanced sample and enhanced genes were re-randomized for each trial.

#### Gene-gene correlation tests

Figure 11 compares gene-gene Pearson correlations computed using the original log transformed data, and using local-leveled log transformed data. To obtain the curves labeled “Local correlation”, first the genes were sorted in order of increasing log-transformed expression level, and 1000 consecutive genes starting with index 12000 were selected. All correlations between these genes were computed, and the resulting distribution is plotted. To obtain the curves denoted “Random correlation”, 1000 genes were randomly selected from among all genes with index above 7000 (low-expressed genes we excluded because they are often unexpressed and have highly irregular statistics). The figure shows that local and random correlations for the original data both show a positive shift which is especially pronounced for the local correlations. This result is an expected consequence of the expression level-dependent block shifts observed in Figures 6 and 7. On the other hand, the local leveled correlations for both distributions are centered at zero, and the distributions are nearly the same.

**Figure 11.**
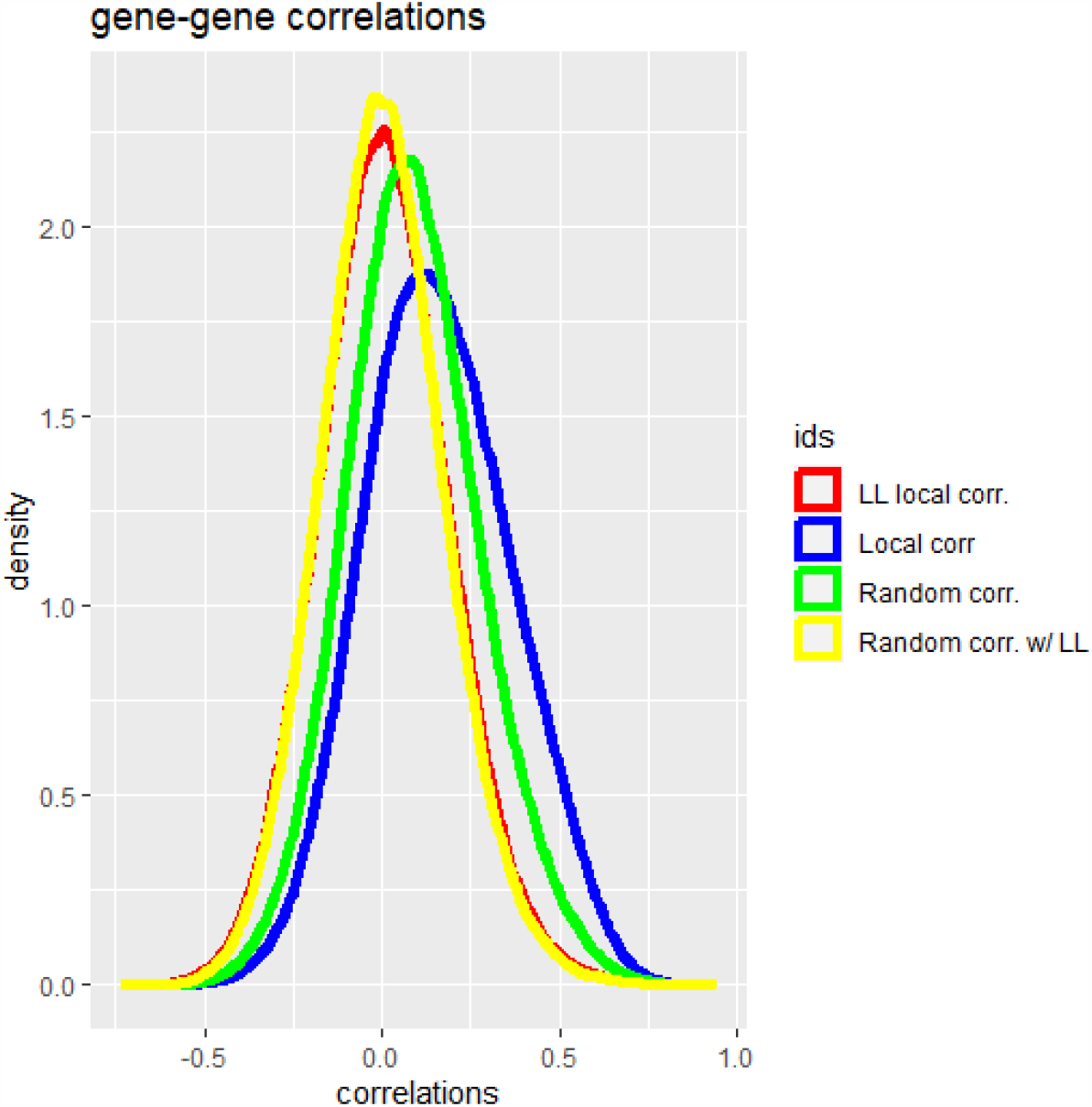
Gene-gene log transformed expression level correlations between intervals of genes with similar expression levels (local corr.) and genes with randomly-selected expression levels (random corr), for original bladder cancer TPM data and local leveled data.

Figure 12 shows Spearman correlations between a particular gene (the androgen receptor gene) and all other genes, for all samples within the TGCA firehose prostate TPM dataset. In this case the original data is used without applying the log transform, while the local leveled data is implemented by first log transforming the data, then local leveling, then applying the inverse transform. The figure shows that the correlations using original TPM data are skewed positive, with a large proportion of values above 0.5. In contrast, the distribution of correlations computed with local leveling is much more symmetric, more sharply peaked and centered near 0. The reduction in correlation indicates that local leveling has corrected for correlations due to the per-patient shifts shown in Figure 6.

**Figure 12.**
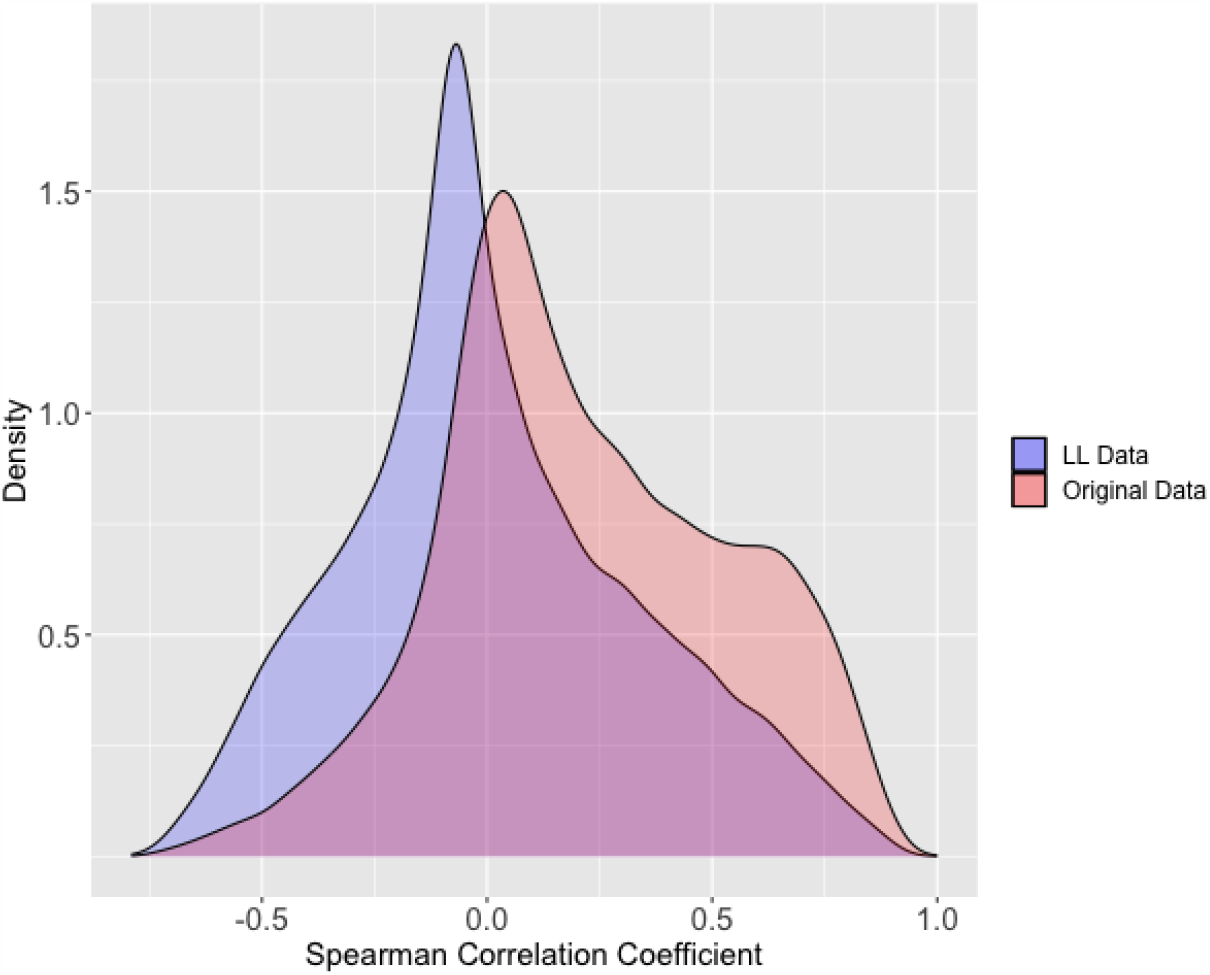
Distribution of Spearman correlation coefficients between androgen receptor (AR) gene and all other genes, for original and for local leveled data. Data is taken from TGCA prostate firehose dataset.

## 6 Discussion

### 6.1 Possible causes of systematic bias

In Section 3 Although we have shown figures only for TPM bladder cancer data from TGCA, similar figures can be generated for all datasets described in Table 1. In Section 5 we showed that these biases do affect tests designed to identify genetic differences in populations and individuals, as well as correlations between genes. We show furthermore that local leveling is partially effective in mitigating the effects of systematic biases.

Until now we have not identified the cause(s) of these systematic biases. One candidate is faulty normalization. But faulty normalization cannot fully explain the phenomenon. Data is normalized by multiplying all expression levels by the same constant. Under log transform, this multiplication translates to a constant shift that is applied equally to all genes. Thus faulty normalization can account for overall shifts in log transformed expression levels between samples, such as shown in the scrambled samples in Figure 6(b). But faulty normalization cannot explain the expression level-dependent shifts in the log transformed samples that are seen in Figure 6(a). We note however, that the vertical shifts in scrambled samples can be eliminated by adjusting the normalization, thus improving the statistical behavior. The shifts in Figure 6(b) are clear evidence that TPM normalization is not accurate, and the shifts can be used to correct the multiplicative factor used for normalization.

Another potential source of systematic bias is errors in the gene length estimates that are used to convert fragment counts to fragment counts per kilobase. This is because the systematic variations also show up in the raw data, even before the conversions are made.

A further possible cause is mapping error. Mapping errors in the context of intrepreting RNA-seq data are well documented in the literature [2]. These inaccuracies arise during the alignment or mapping step of RNA-sequence analysis. This step involves aligning the short sequence with a reference genome to determine where it comes from. Due to many factors (e.g. repetitive regions, alternative splicing, sequencing errors, etc.), reads may be incorrectly assigned, leading to biases in the gene expression levels. Researcher have used different statistical methods to deal with these biases [2]. However, it is unclear why mapping error would be related to gene expression level. This is a topic for further investigation.

In sum, we have so far been unable to identify a cause for the observed biases. We cannot rule out the possibility that what appears to be measurement bias may in fact be of biological origin.

### 6.2 Implications of local leveling on population level data

A current need in the field of cancer is to interpret how genes can drive tumorigenic processes and the pathogenesis of human tumors. As discussed, RNA-seq or other forms of abundance level data are now collected at the population level across hundreds of patients in the research setting, and even millions of patients in the industry setting. Because of this, a specific need is to consider batch and inter patient normalization processes when abundance level data is acquired, which may include biases or artifacts that are due to the variations in collection process or differences in sequencing platform. These processes are areas of active research and development. Local leveling acts as once such algorithm, as it considers expression levels of genes from the entire population of interest when making adjustments. Unlike other normalizations, the local leveling algorithm takes into account both gene-to-gene and patient-to-patient variations. TPM normalization or log transform apply the same transform to all genes on a per-sample basis. Such transformations will reduce spurious variations in gene expression levels, but do not address gene-to-gene differences in these spurious variations. In contrast, Z-score normalization applies the same linear transformation to all samples, where the transformation is determined independently for each gene. As a result, the Z-score transformation has no effect on gene-gene correlations. (Note that in our application of local leveling we also apply Z-score normalization, but only *after* local leveling has been applied.)

In this study we have applied local leveling only to RNA-seq data. It is possible that the same technique may be applied to reduce biases in protein abundance level estimates obtained from mass spectrometry data. This is a topic for future investigation.

#### 6.2.1 Impact on biological functions and signaling pathways

## 7 Implications and conclusion

Data processing often introduces distortions and biases. Methods must be devised to correct for these. In this paper, local leveling is proposed and validated as a method for correcting biases in RNA-seq data. These corrections may significantly effect various biological analyses. Some possible consequences are described as follows.

In this study, we have demonstrated that local leveling has a global impact on gene-gene correlations. In particular local leveling shifts the mean of the distribution of the correlations towards zero. Conventionally, gene-gene correlations across all the samples are often used as a measure of the degree of similarity between two or more genes.

Differential expression analyses through bioinformatics tools such as DEseq2 [5] stratify patients into groups based on certain clinical or experimental annotations, and then use statistical models to identify specific genes that exhibit different expression levels between groups. The information is commonly used to define potential biomarkers, or to nominate downstream experimental strategies. Local leveling will impact differential expression analysis, and we predict that this will nominate distinct gene sets that could inform of novel roles for certain genes.

Local leveling may also impact gene set enrichment tools such as GSEA [7], that evaluate whether specific biological pathways are active or inactive in samples. Like differential expression analyses, these tools examine whether an a priori nominated gene set exhibits concordant differential expression. Many of these gene sets are defined based on optimized conditions in cell models, which are generally more synchronous and homogeneous as compared to biological samples. In this regard, the genes sets may exhibit much more pronounced activity and less variation. However, gene set enrichment approaches are also applied on clinical samples. Unlike cell models, these are subject to many distinct collection and processing protocols on heterogeneous cell types or tumor cells. Each of these factors may drive sub-optimal expression of genes. For these reasons, local leveling may have a greater impact on studying gene sets or biological pathways in clinical samples. Altogether, applying local leveling on more clinical specimens may provide novel understanding of gene-gene interactions and biological pathways.

Further investigations are being pursued into characterizing the observed biases more precisely, with a view to identifying possible causes. In particular, there may be other factors besides gene expression level that are associated with systematic biases. Identifying and compensating for these factors can lead to better bias correction procedures, thus producing data that is more representative of underlying biological conditions.

## Notes

### Competing Interest Statement

The authors have declared no competing interest.

